# Degraded neural representations foreshadow failures of sustained attention

**DOI:** 10.64898/2026.09.21.753248

**Authors:** Anna Corriveau, Dongfang Tian, Matthieu Chidharom, Edward K. Vogel, Monica D. Rosenberg

## Abstract

Attention prioritizes relevant information from the environment, but does not do so consistently over time. Instead, we sometimes experience poor attentional states which are characterized by a higher degree of performance failures, or “lapses.” The source of these lapses—whether they arise from attentional disengagement, failures of attentional selection, or simply failures of response control—is not clear. This study tests whether the strength of neural representations differs as a function of both attentional selection and modulation over time. We recorded electroencephalography (EEG) data while participants performed a sustained attention task requiring the selection of task-relevant items and withholding of responses to rare targets.Results show that relevant items receive prioritized processing in brain signals, as measured with EEG decoding. Yet, their neural representational fidelity weakens under poor sustained attentional states, foreshadowing upcoming behavioral failures. These findings support a neural basis of disengagement during poor attentional states, suggesting that lapses may arise from a decoupling of brain activity from task processing. Finally, pre-item representational strength predicted memory success for targets, suggesting that neural fidelity indexes an encoding-ready state. Results suggest that the quality of neural representations is affected by ongoing attention and plays a key role in processing and memory.

**Significance Statement:** Human attention is dynamic, leading at times to error- or “lapse”-prone states. Yet, there is debate about whether attentional lapses are driven by disengagement from relevant information or simply reflect failures to respond appropriately to that information. We show that neural task representations degrade before lapses, supporting the disengagement account. Representational fidelity dynamics are not explained by irrelevant-item distraction and predict subsequent memory, indicating that momentary neural processing indexes both an engaged and encoding-ready state. Attention lapses and resulting encoding failures therefore reflect disruptions to information processing rather than response control.

## Introduction

Despite our best intentions, humans struggle to maintain the focus of attention on task-relevant goals for extended periods of time, instead fluctuating between periods of in-the-zone and error-prone, out-of-the-zone processing. Yet, it is not clear whether sustained attentional lapses stem from disengagement from task processing or whether they are simply a failure of response control. Using electroencephalographic (EEG) recordings, we monitored representational strength of task-relevant and task-irrelevant information under fluctuating sustained attention defined by response errors. Results reveal that relevant item representations, in addition to receiving prolonged priority over irrelevant information, are stronger under better sustained attention, supporting attentional disengagement as a driver of lapses. We further extend these findings to better understand how attention-driven representational changes impact memory, providing novel insights into how dynamic attentional states shape both behavior and cognition.

A growing body of work demonstrates that attentional selection impacts the strength of neural representations. Work across species and modalities demonstrates that attention strengthens the fidelity of neural signals ^1^ and representations of selected items ^2–5^ and features ^6,7^. Changing task-relevant goals leads to widespread shifts in neural geometry and cortical tuning ^8–13^ as well as patterns of hippocampal activity ^14^, suggesting that neural activity patterns adapt to accommodate attentional selection. In combination, this work indicates that selective attention reliably increases the availability of information in neural signals, strengthening representations of relevant items.

However, the effects of sustained attention dynamics on neural representations are not well established. Fluctuations in attentional state—here used synonymously with sustained attention—occur dynamically across moments ^15^. While sustained attention can be tracked with patterns of brain activity ^16–20^, it is not clear whether these states reflect differences in task-specific processing or a condition unrelated to external input (e.g., arousal). Attentional fluctuations are frequently measured using continuous performance tasks, which require behavioral responses (e.g., button presses) to frequent items and the withholding of responses to rare, infrequent items ^21,22^. Failure to withhold a response on rare trials reflects a sustained attentional lapse and suggests that the focus of one’s attention is somewhere other than the immediate task goals. However, the basis of sustained attentional lapses is unclear. One possibility is that poor sustained attentional states are characterized by attentional disengagement, or decoupling between the brain and stimulus information, with lapses resulting from a failure to adequately detect a rare target ^21,23–25^. Supporting this view, stimulus-evoked brain activity changes under differing attentional states ^26,27^ and aligns across individuals under high sustained attention ^28^. Additionally, networks sensitive to sustained attention lapses predict attention under entirely different contexts, including decoupling during internally-generated thought ^29^, tracking subjective fluctuations in narrative engagement ^30,31^ and selectively reconfiguring under administration of drugs that impact attention ^32–35^.

An alternative explanation, however, is that lapses instead reflect a shift in response bias and failure of motor inhibition when the prepotent response needs to change ^36^. Evidence for this view comes from findings that frequent responding leads to more errors on rare trials than tasks requiring infrequent responses ^37,38^, that manipulating task instructions or demands (e.g., delaying responses) reduces lapses ^39–42^, and that individuals are often aware of errors after they occur ^43^. While it is possible that attentional disengagement and failures of response control may both contribute to sustained attentional lapses ^44^, some proponents of the response control view maintain that these results preclude attentional disengagement during tasks involving frequent responding from contributing to sustained attention errors at all ^36,45^.

Critically, these views make separate predictions for how fluctuations in sustained attention should impact task-related neural activity patterns. If lapses are to some extent due to disengagement from the task, we should expect differences in the representational strength of relevant items in brain signals during poor attentional states. Further, if lapses are caused by misdirected attention towards irrelevant information, irrelevant representations may strengthen prior to lapses. If, on the other hand, errors are driven purely by failures of motor inhibition due to frequent responding, neural representations should not signal differences between good and lapsing sustained attention. In the current study, we utilize EEG decoding to examine whether trial-wise representations portend lapses in sustained attention, and how representations differ as a function of task relevance.

In addition to changes in processing, both selective and sustained attention impact how stimuli are subsequently remembered ^46,47^. Task-relevant stimulus features are better remembered ^14,48–50^, as are stimuli presented in selected locations in space ^51,52^. Information presented during better sustained attentional states, as indexed by online behavioral measures, also show superior memory performance ^53–55^. Interestingly, the impacts of selective and sustained attention on recognition memory performance appear to be dissociable ^48,49,56^ such that both types of attention uniquely contribute to successful memory judgements. For example, Corriveau & Chao et al. ^48^ found that, atop a robust effect of selective attention (task-relevance) on memory judgements, both relevant and irrelevant images presented during better sustained attentional states were better-remembered. These results, in combination with related work ^49^, contradict the intuition of an inherent tradeoff between processing for relevant and irrelevant information and necessitate investigation into the unique impacts of selective and sustained attention on mnemonic processes.

The primary aim of the present study is to examine how the engagement of attention—both prioritization and dynamic states—impacts neural representations of visual stimuli. Weakened representations under poor sustained attentional states would support attentional disengagement as a driver of lapses, beyond simply failures of response inhibition. Participants completed a modified continuous performance task in which they selectively attended specific stimuli within a spatially-consistent visual display. We examined how the availability of stimulus information in neural signals varied as a function of selectivity (task-relevance) and sustained attentional state using EEG decoding. In line with previous work ^3^, we find that selective attention strengthens the representational fidelity of task-relevant information. Further, we show that sustained attention errors are foreshadowed by weakened stimulus representations, suggesting that lapses may be driven by decoupling between the brain and task-relevant processing. No corresponding increase in irrelevant representations prior to errors suggests that lapses are not simply a failure of attentional selection. Finally, we find that item-level memory is predicted by preceding representational strength, suggesting that the fidelity of neural representations reflects both processing and encoding success.

## Results

Participants (N=25, mean age=22.24 years, SD=2.93 years) performed a continuous performance task (CPT) which measures behavioral lapses in sustained attention while EEG recorded electrical activity across the scalp (**Figure 1**). Stimuli were images superimposed in the center of Gabor patches. To manipulate the focus of selective attention, participants were instructed at the start of each task run to make a judgement about either the category of images or the orientation of Gabor patches, with frequent trials equally drawn from one of two categories (images of food or vehicles, horizontal or vertical Gabor patches) and infrequent trials drawn from a third rare category (images of animals, diagonally-oriented Gabor patches). Participants were tasked with responding to frequent-category stimuli (90% of trials) while withholding responses to infrequent-category stimuli (10% of trials). Sustained attention performance was measured discretely using accuracy on infrequent trials, where correct omissions indicate good sustained attention and errors of commission indicate an attentional lapse. Lapse rates (Image-relevant runs: mean=.226, SD=.098; Gabor-relevant runs: mean=.307, SD=.136) were in line with rates reported previously ^57,58^ and omission rates were low, as expected (Image-relevant run: mean=6.63*10^-3^, SD=9.21*10^-3^; Gabor-relevant runs: mean=3.22*10^-3^, SD=4.15*10^-3^). We also tracked continuous fluctuations in sustained attentional state using responses on correct, frequent trials by calculating time courses of response speed and variance, which have been shown to predict sustained attention errors ^22,48,49,54,55,57,59^. Average response times for correct, frequent trials were comparable between Image-relevant (mean RT=.462s, SD=.046s) and Gabor-relevant runs (mean RT=.460s, SD=.047s). See Materials and Methods for complete task descriptions.

**Figure 1.**
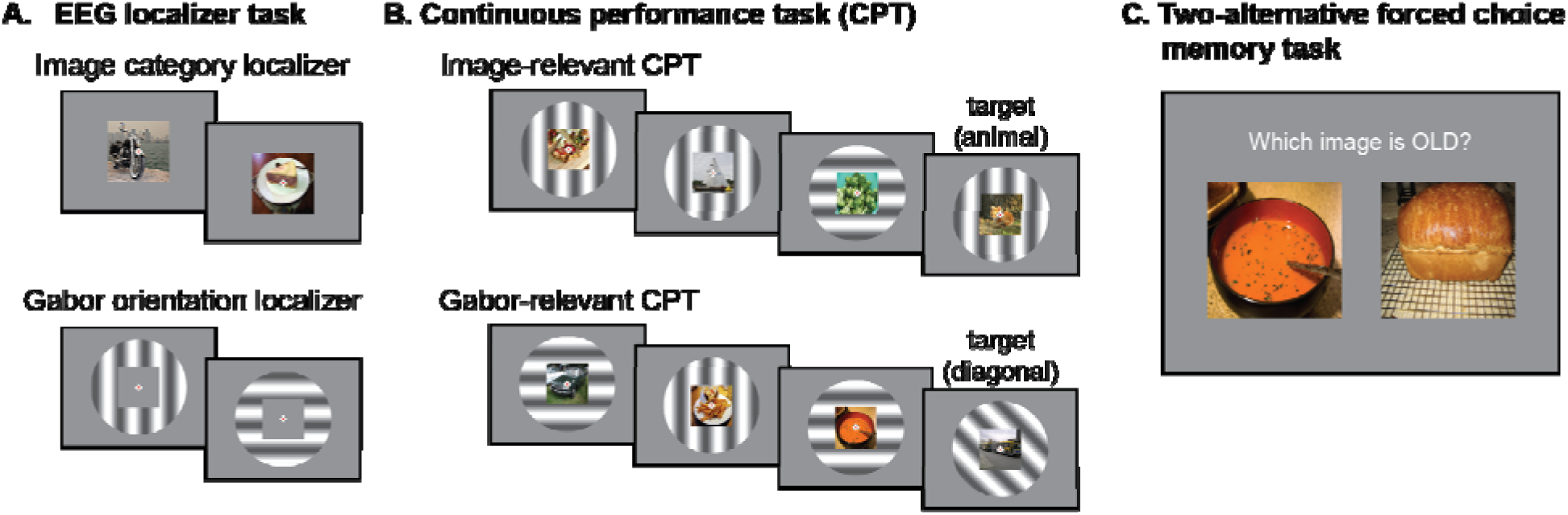
*Experimental procedure*. (A) Participants completed three tasks as part of the experimental procedure. Task images are visualizations and are not presented to scale. The EEG localizer task included six alternating blocks of images (3 blocks) and Gabor patches (3 blocks). Participants detected a color change at fixation (red to green) which occurred on 10% of trials. Stimuli were presented for 800 ms with a 200ms inter-trial interval (ITI). Blocks included 600 trials each. (B) The continuous performance task featured stimuli composed of images superimposed on Gabor patches. Participants were assigned to make a judgement of image category (image-relevant runs) or Gabor orientation (Gabor-relevant runs), pressing to frequent-category stimuli (food and vehicle images; horizontal and vertical Gabor patches) and withholding responses to rare targets (animal images; diagonal Gabors). Stimuli were presented for 800 ms with a 500 ms “blink” period and a 200 ms ITI. Participants completed four CPT blocks–two image-relevant blocks and two Gabor-relevant blocks in alternating order. (C) Finally, participants completed a surprise two-alternative forced choice memory task for images presented during CPT blocks. Participants were tasked with identifying the old image and provided a confidence rating (definitely or maybe) with a key mapping (z=definitely left; x=maybe left; n=maybe right; m=definitely right). This key mapping remained on the screen for all trials but is not depicted due to limited space. Participants completed 300 self-paced memory trials (180 relevant images, 120 irrelevant images).

The aim of the current study was to test whether the classification of frequent-category stimuli (food vs. vehicle images; vertical vs. horizontal Gabor patches) from EEG signals differs as a function of attention, which would indicate differences in the amount of stimulus information available in brain signals under different attentional conditions. The task designed allowed us to compare unique contributions of both selective and sustained attention to differences in EEG classification performance by evaluating performance in task runs where stimuli were relevant vs. irrelevant (selective attention), as well as when participants were in periods of better vs. worse sustained attentional states, defined by trial-wise performance and response times. To remove confounding factors of motor responses to EEG signals, within-subject linear-discriminant EEG classifiers were trained on separate task runs (EEG localizer task, see Methods) during which participants made a response only on rare, attention-check trials which were not included in classifier training. This study was preregistered at https://osf.io/h9r4e/overview. We report preregistered and exploratory analyses as appropriate throughout the manuscript.

### Selective attention prolongs maintenance of relevant information

Task performance (sensitivity, *A’*) was high in both the EEG localizer task used for classifier training, where *A’*=.5 represents chance-level performance (image runs: mean *A’*=.993, SD=7.34*10^-3^, Gabor runs: mean *A’*=.993, SD=8.27*10^-3^) and the CPT used for classifier testing (image-relevant runs: mean *A’*=.941, SD=.027; Gabor-relevant runs: mean *A’*=.921, SD=.036), suggesting that participants maintained attention well. Participants also performed consistently on the EEG localizer task (*r=*.795, p<.001) and the CPT (*r=*.728, p<.001), indicating reliable individual differences in attentional performance.

Because both image and Gabor patch stimuli were presented on all trials but varied by task-relevance, we were able to test whether the availability of stimulus information in EEG signals as measured with classification performance varied with the focus of selective attention. Our first registered analysis evaluated the time course of EEG classification performance throughout the trial duration to determine whether and when task relevance impacted how information was reflected in EEG activity. We hypothesized that neural representations of task-relevant stimuli would be more reliably reflected in EEG signals, enabling more successful decoding. In line with our hypothesis, classification of task-relevant images (mean AUC=.554, SD=.022) from EEG signals was more accurate than classification of task-irrelevant images (mean AUC=.530, SD=.017) on average throughout the trial (t(24)=5.80, p<.001, BF_10_=3671.50; **Figure 2A**). Average trial-wise decoding performance was above-chance for both relevant (t(24)=11.74, p<.001, BF_10_=9.13*10^8^, one-sided) and irrelevant images (t(24)=8.67, p<.001, BF_10_ = 3.39*10^6^, one-sided). Examining the time course of image decoding, differences emerged between 300ms through 600ms after stimulus onset, with task-irrelevant image decoding dropping more rapidly during this time, suggesting that image category information was not as well maintained in EEG activity. The initial rise in decoding performance did not differ by task-relevance, indicating consistent timing of image category classification regardless of task-relevance.

**Figure 2.**
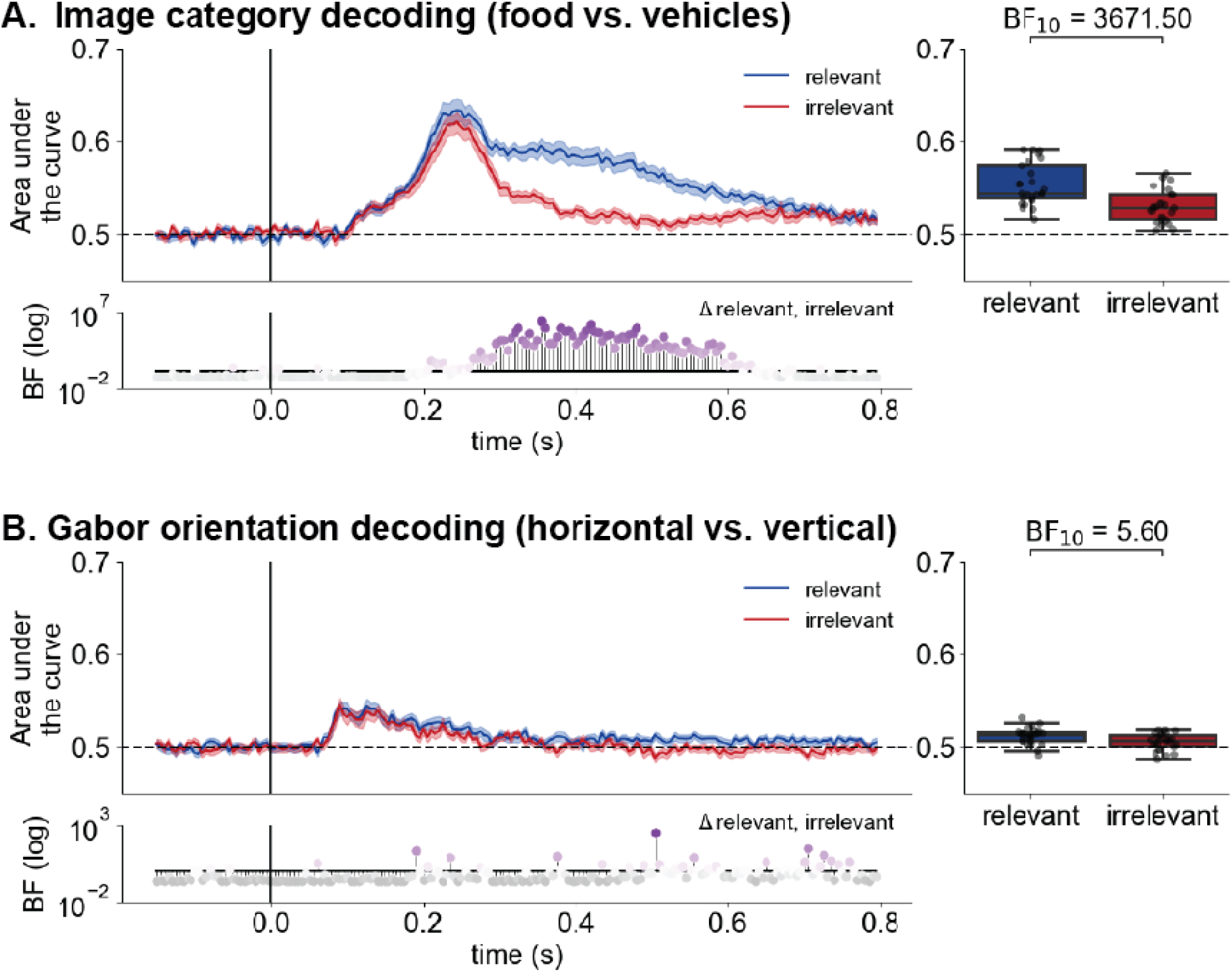
*Stronger decoding of task-relevant stimuli.* The classification of (A) image categories and (B) Gabor orientations during a continuous performance task is higher when stimuli are task-relevant. Bayes factors (BFs) reflect evidence of differences between task-relevant and task-irrelevant time courses across participants. BFs greater than one are shown in color and indicate evidence of a difference. Box-and-whisker plots reflect subject-wise mean area under the curve across the trial (0-800 ms after onset).

Performance for decoding the orientation of Gabor patches was low on average for relevant (mean AUC=.512, SD=8.93*10^-3^) and irrelevant Gabor patches (mean AUC=.506, SD=8.76*10^-3^), although trial-average decoding was above-chance for both relevant (t(24)=6.70, p<.001, BF_10_=5.56*10^4^, one-sided) and irrelevant Gabor patches (t(24)=3.13, p=2.30*10^-3^, BF_10_=18.51, one-sided). We observed moderate evidence for an effect of selective attention such that relevant orientation decoding was higher than irrelevant decoding on average (t(24)=2.88, p=8.29*10^-3^, BF_10_=5.60; **Figure 2B**). Examining the time course of classification across the trial, task-relevant orientations remained higher late in the trial, suggesting that information about Gabor orientations were maintained more strongly throughout the trial’s duration when Gabor patches were the relevant stimulus. In sum, the attentional selection afforded to task-relevant stimuli leads to prolonged maintenance of stimulus information in EEG signals as indicated by stronger classification performance.

To determine whether the attentional selection boosted the decodability of relevant stimuli or interfered with the decodability of irrelevant stimuli, we compared classification performance during CPT to performance during localizer blocks, with which the classifiers were trained. Task-irrelevant stimulus decoding during the CPT was lower than decoding during the localizer task, suggesting that task-irrelevant stimulus information is more suppressed during the CPT (**Figure S1**).

Finally, we examined whether decoding performance was driven by eye movements. We retrained classifiers to decode image categories and Gabor orientations using eye gaze and pupil data (see **Methods**). The time course of decoding from eye-based data is distinct from that of EEG classifiers (**Figure S2**). Further, predictions from EEG- and eye-based classifiers are not differentially correlated between relevant and irrelevant predictions, suggesting that eye-related features do not drive the differences observed as a function of task relevance (**Figure S2**).

### Sustained attention lapses are preceded by degraded representations

Having established an effect of attentional selection, i.e., stronger decoding performance for task-relevant information, we next tested whether fluctuations in sustained attention impacted trial-wise decoding performance. During tasks like the CPT in which participants must inhibit responses to infrequent targets, erroneous presses to target stimuli are often interpreted as sustained attentional lapses, characterized by disengagement from task processing ^21^. It is possible, however, that lapses are not due to poor attention *per se*, but rather a failure of motor processes to inhibit a response ^36^. Here, we tested whether CPT target performance is foreshadowed by differences in representational strength in EEG signals in the three-trial window preceding target presentation. Weakened decoding strength prior to an attentional lapse itself would suggest that errors are characterized by disengagement from task-related processing and are not simply a byproduct of response control failure. A similar trailing window approach has previously been shown to predict upcoming sustained attention lapses both behaviorally ^48,49,53–55,60,61^ and neurally ^17^. Due to poor decoding performance of Gabor orientations as predicted in our preregistration, main results focus on the time course of image category decoding. We report Gabor orientation decoding results in the supplement (**Figure S3**).

Relevant images presented prior to an erroneous response to a rare target (mean AUC=.529, SD=.047) showed worse decoding relative to those preceding a correctly-withheld response (mean AUC=.561, SD=.028), suggesting that lapses are portended by weakened brain representations (t(24)=4.04, p<.001, BF_10_=152.2; **Figure 3A**). This difference began ∼300ms after onset and persisted for the remainder of the trial. These results indicate that upcoming sustained attention performance is predicted by the availability of stimulus information in EEG signals such that lapses are foreshadowed by degraded representations of relevant content.

**Figure 3.**
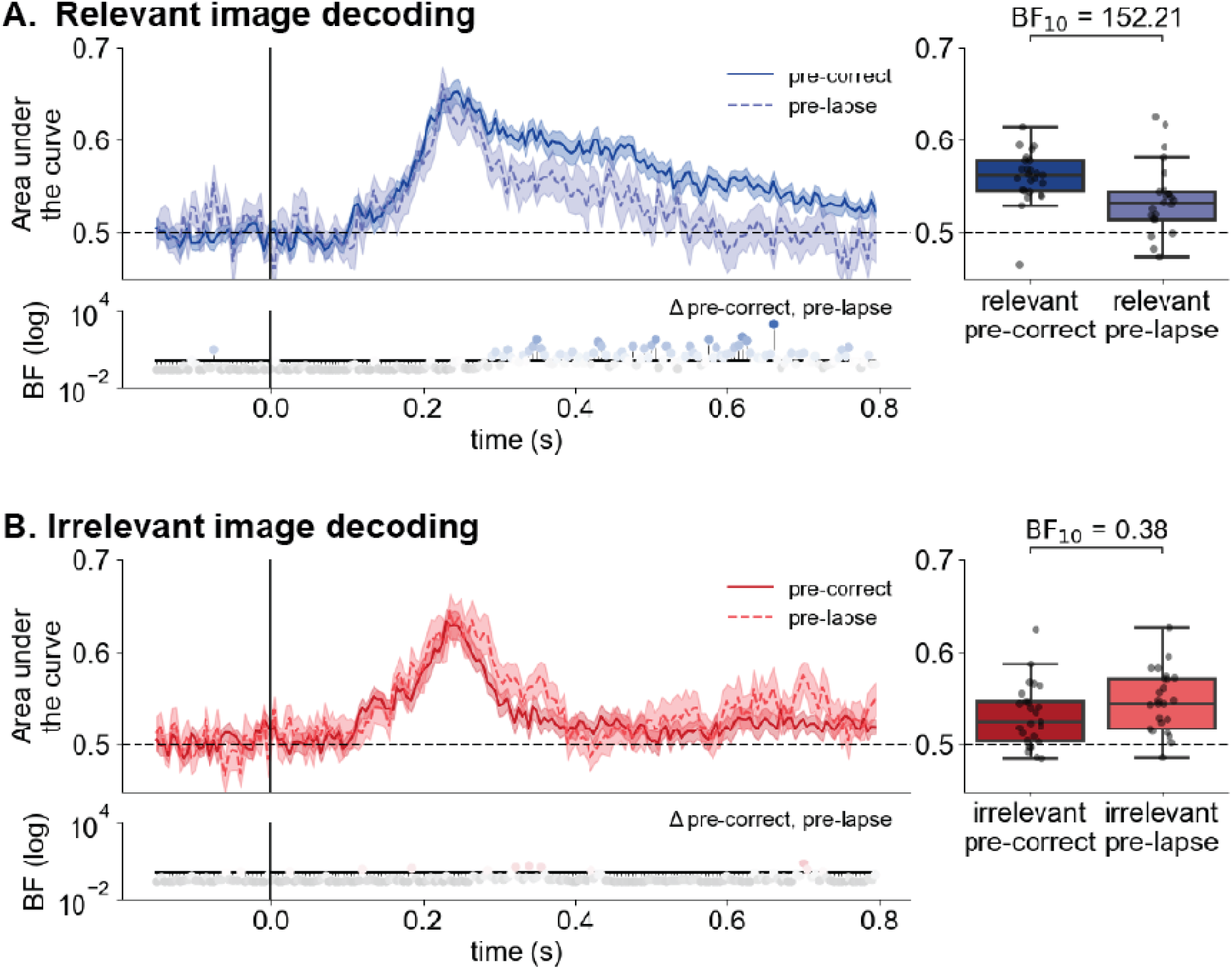
*Weak neural representations of relevant information prior to sustained attentional lapses.* Decoding performance was poorer for (A) relevant stimuli preceding sustained attentional lapses suggesting that neural representations are weaker prior to poor sustained attention performance. (B) Irrelevant image decoding did not distinguish lapsing vs. correct sustained attention performance on average. Box-and-whisker plots reflect subject-wise mean area under the curve across the trial (0-800 ms after onset).

A Bayesian mixed effects model predicting decoding performance from target trial performance and trial number found that decoding performance is reliably related to target trial performance (mean effect=.126, 89%CI[.051, .200]) despite waning in later trials (mean effect=-.046, 89%CI[-.077, -.015]), suggesting that degraded representations are due to lapses themselves and not weakened signal over time. Additionally, better decoding performance prior to correctly-withheld responses is not likely due to a greater number of trials being tested. While participants indeed were more likely to correctly withhold responses (mean trials=167.72, SD=43.23) than lapse (mean trials=47.0, SD=19.44), robust differences were maintained when the number of test trials was equal across conditions (**Figure S4**).

We did not observe differences in the average decoding of irrelevant image categories as a function of sustained attentional performance (mean AUC_correct_=.532, SD_correct_=.033; mean AUC_lapse_=.543, SD_lapse_=.040; t(24)=-1.15, p=.263, BF_10_=0.38; **Figure 3B**). Across the time course, small differences emerge in irrelevant image decoding performance, with pre-lapse decoding briefly and inconsistently out performing performance under better sustained attentional states. However, the lack of reliable differences throughout the trial suggests that irrelevant stimulus representations do not predict upcoming sustained attention failures under the conditions tested here and, further, that lapses are not simply due to a shift in attention to irrelevant information. A Bayesian linear model predicting average pre-trial decoding performance from task relevance, target-trial performance (correct vs. lapse), and their interaction confirms that relevance (mean effect=.029, 89%CI[.011, .047]) and the interaction between relevance and target performance (mean effect=-.043, 89%CI[-.068, -.018]) predicts decoding performance, such that higher decoding as a function of target performance occurs only when images are task-relevant. Target performance alone did not significantly predict decoding success (mean effect=.012, 89%CI[-6.11*10^-3^, .029]). Taken together, results suggest that the processing of relevant items specifically degrades prior to sustained attentional lapses.

Are differences in decoding performance preceding better vs. worse sustained attention performance driven by differential evoked EEG activity under different states? We next examined whether the onset of images prior to correctly-withheld responses elicited different levels of raw EEG activity than those prior to sustained attentional lapses by examining the differences in evoked response potentials (ERPs) across channels. We observed little to no evidence of ERP differences across the scalp preceding correct vs. lapsing sustained attention performance (**Figure S5**).

Finally, as an exploratory analysis, we tested whether posterior alpha power, thought to be involved in the inhibition of irrelevant information ^62^, differed between trials preceding correctly-withheld responses (good sustained attention) and attentional lapses by examining the differences in the time-frequency decomposition between conditions. Low frequency power including posterior alpha power (8-12 Hz), was greater on trials preceding sustained attentional lapses around the time of stimulus onset (**Figure S6**) as well as prior to trial offset, possibly indicating greater reliance on top-down inhibition during poor attention, analogous to observations of greater dorsal attention network activity during out-of-the-zone attentional states ^15^. This suggests that differential sustained attentional performance is predicted by differences in oscillatory power in low frequency bands and these differences are most pronounced at key moments in a trial (e.g., stimulus onset and offset). However, while alpha power may impact decoding performance, a Bayesian mixed effects model predicting decoding performance from both target performance and trial-averaged alpha power confirm that correct target performance predicts more successful decoding (mean effect=.079, 89%CI[.027, .132]) when accounting for alpha (mean effect=-8.52*10^-3^, 89%CI[-.030, .013]). This suggests that, while alpha power differs with attentional performance, it does not explain weakened representational strength of images during lapses.

### Individual differences in selective processing tradeoffs

Selective attention is classically thought of as a spotlight, such that relevant information in the focus of attention is prioritized while irrelevant information is suppressed ^63,64^. By extension, it may be supposed that lapses in attention reflectreflect selective attention failuress, resulting in increased processing of irrelevant information. Under this view, we would expect neural representational strength to trade-off over time, such that when representations of relevant information are high, irrelevant representations should be poor, and vice versa. However, previous work finds that better processing of relevant stimuli during the CPT does not come at the cost of irrelevant stimulus processing in adults ^48,49,65^. Therefore, in an exploratory analysis, we leveraged trial-level classifier predictions to test whether classification performance for relevant and irrelevant stimuli within the same trial fluctuate in sync or in antiphase across trials. We correlated decoding performance for relevant and irrelevant stimuli across all trials within a subject. A positive correlation would suggest simultaneous processing of relevant and irrelevant stimuli; a negative correlation would suggest a tradeoff such that increased processing of relevant stimuli is accompanied by decreased processing for irrelevant stimuli and vice versa.

On average, we did not find evidence for either a simultaneous boost nor a tradeoff in image-relevant (mean *r_relevant, irrelevant_*=-.056, SD=.332; One-sample t-test: t(24)=-.822, p=.420, BF_10_=.286) and Gabor-relevant runs (mean *r_relevant, irrelevant_*=-.045, SD=.316; One-sample t-test: t(24)=-.694, p=.495, BF_10_=.262). This result adds to growing evidence that, on average, processing for relevant information does not increase at the expense of irrelevant processing, or vice versa ^48,49,65^. However, we observed strong consistency across participants (*r*=.964, p<.001) suggesting that the level of processing tradeoffs is reliable within an individual. In other words, individuals who show simultaneous boosts in processing for relevant and irrelevant information do so regardless of the relevant stimulus category.

### Response time measures predict attention lapses but not representational differences

Because participants were tasked with responding to the vast majority (90%) of CPT trials and made very few omission errors (mean_image-relevant_=.67%, SD=.92% of trials; mean_Gabor-relevant_=.32%, SD=.42% of trials), we are able to monitor continuous changes in sustained attentional states through measures related to response time. Specifically, previous work demonstrates that individuals are more prone to sustained attentional lapses during states quantified with both speeded ^53,55^ and highly variable responding ^22,57^. In the current study, Bayesian mixed effects modeling with effects of response speed, variance, their interaction, and block type (image-relevant or Gabor-relevant) found evidence that behavioral responding predicted performance on infrequent sustained attentional probes during the CPT. Specifically, correctly-withheld responses were preceded by slower (mean effect=0.515, 89%CI[0.448, 0.584]) and less variable pressing (mean effect=-0.181, 89%CI[-0.239, -0.122]). We also found evidence of an interaction between response speed and variance (mean effect=-0.053, 89%CI[-0.077, -0.031]) such that slow, stable responses were more likely to precede correct responses than slow but variable responses. Finally, participants made fewer attentional lapses in image-relevant CPT blocks relative to Gabor-relevant blocks (mean effect=0.473, 89%CI[0.373, 0.575]). Results replicate previous work suggesting that behavioral indices of speed and variability are uniquely related to trial-level sustained attentional performance.

We predicted that differences in decoding performance would emerge as a function of sustained attentional state as measured with response speed and variance. While our preregistered analyses described performing median splits based on response speed and variance, we opted to analyze the most upper and lower quartiles of trials (i.e., those occurring under the most extreme sustained attentional states as determined by response measures) to provide the most power for detecting a difference. Even when analyzing trials in the best and worst sustained attentional states as indicated by response speed (Relevant images: mean AUC_high_=.553, SD_high_=.027; mean AUC_low_=.556, SD_low_=.029; t(24)=-.493, p=.626, BF_10_=.236; Irrelevant images: mean AUC_high_=.531, SD_high_=.021; mean AUC_low_=.528, SD_low_=.023; t(24)=.550, p=.587, BF_10_=.242) and variance (Relevant images: mean AUC_high_=.555, SD_high_=.028; mean AUC_low_=.550, SD_low_=.027; t(24)=.871, p=.392, BF_10_=.297; Irrelevant images: mean AUC_high_=.532, SD_high_=.024; mean AUC_low_=.529, SD_low_=.024; t(24)=.549, p=.588, BF_10_=.242), we did not observe differences in trial-averaged image category decoding for either relevant or irrelevant images (**Figure 4**). Timepoint-by-timepoint differences were brief and inconsistent, suggesting that the current study design lacked the ability to detect differences in stimulus representations as a function of continuous attentional state changes. We similarly observed no differences in Gabor orientation decoding as a function of response-indexed sustained attention across the trial (**Figure S7**).

**Figure 4.**
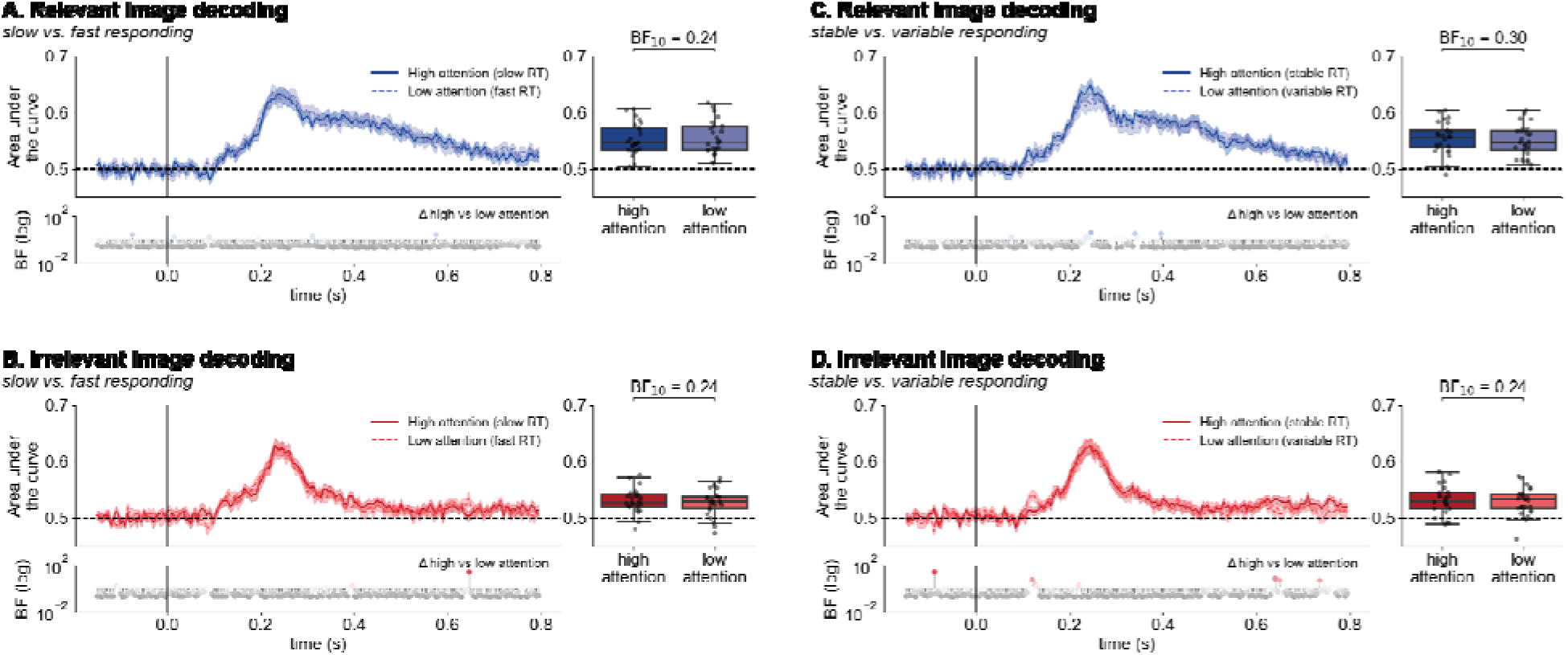
*Response time measures of sustained attention do not impact decoding.* Decoding performance of relevant and irrelevant images did not reliably differ as a function of sustained attentional states as determined by response time (RT) speed (A-B) nor variance (C-D). Box-and-whisker plots reflect subject-wise mean area under the curve across the trial (0-800 ms after onset).

In combination with previous results showing weakened decoding success prior to attentional lapses, these results suggest that differences in response times under better and worse sustained attentional states are unlikely to explain the weakened decoding observed. To confirm this, we fit a Bayesian mixed effects model predicting decoding performance from target trial performance with additional effects of response speed, variance, and their interaction. While target trial performance remained a reliable predictor of decoding performance (mean effect=.132, 89% CI[.057, .208]), response speed (mean effect=.029, 89%CI[-.013, .071]), variance (mean effect=.038, 89% CI[9.60*10^-4^, .071]), and their interaction (mean effect=- 6.57*10^-3^, 89%CI [-.025, .011]) were marginally or unrelated to decoding success. Therefore it is unlikely that response characteristics contribute to differences in decoding performance.

### Stronger stimulus representations predict successful memory encoding

Attention impacts what we remember ^46,47^. In the current study, we were interested in whether attention-related measures affected which stimuli were subsequently remembered, in addition to features such as stimulus memorability and false-alarmability, which are known predictors of memory ^66,67^. To test memory in the current study, participants performed a two-alternative forced choice memory task for a subset of images presented during CPT blocks.

Our first exploratory analysis tested what confluence of features impact image memory. We constructed a combination model predicting confident image memory from measures of selective attention (task-relevance), stimulus frequency, and response time predictors of sustained attention. We included additional predictors expected to be related to image memory: memorability, paired lure false-alarmability, and trial number, as well as a subject-level random intercept. We report the posterior distribution mean of estimated effects, where distributions whose 89% credible interval do not include zero are considered evidence of an effect. We also repeated this analysis predicting memory for both confident and unconfident memory judgements, i.e., judgements in which participants correctly selected the old stimulus regardless of confidence report. Results from this model are reported in the supplement (**Table S1**).

When predicting confident memory judgements for all stimuli tested, we observed strong evidence for effects of stimulus memorability (mean= 0.291, 89%CI[0.237, 0.345]), paired lure false-alarmability (mean=-0.141, 89%CI[-0.192, -0.091]), and memory trial number (mean=-0.402, 89%CI[-0.495, -0.309]), suggesting that those stimuli with higher memorability scores, those paired with lures with lower false-alarmability scores, and those presented earlier during the memory task were likely to be remembered. In addition, we observed infrequent targets (animal images) were better remembered than frequent-category images (mean=1.34, 89%CI[1.212, 1.459]), in line with work showing better memory for rare stimuli ^68^. Effects of task-relevance (mean=0.082, 89%CI[-0.121, 0.288]), RT speed (mean=0.064, 89%CI[-0.0044, 0.132]), and RT variance (mean=-0.028, 89%CI[-0.089, 0.033]) were in the expected direction, but the credible intervals of these posterior distributions intersected zero. We saw no evidence of the interaction between RT speed and variance in relation to confident memory judgements (mean=-0.012, 89%CI[-0.043, 0.016]). In sum, both stimulus-related and task-related features impacted successful image recognition.

Finally, following our finding that decoding performance predicted upcoming lapses in the CPT, we were further curious as to whether stronger decoding also foreshadowed successful memory for infrequent target images. To test this question, we analyzed decoding performance, i.e., the classifier-predicted probability that a stimulus belonged to its true image category, in the three-trial window preceding the presentation of an infrequent stimulus. We fit a Bayesian mixed effects logistic regression model predicting confident target memory from effects of decoding performance averaged across the trial, as well as memorability, paired false-alarmability, and memory task trial number. We also included a random intercept of subject. This analysis was proposed as an exploratory analysis in our preregistration. Again, we report results from a model predicting both confident and non-confident memory in the supplement (**Table S2**).

As expected, target memory was predicted by image-level memorability (mean=.211, 89%CI[.115, .308]), the likelihood of the paired lure to elicit false alarms (mean=-.277, 89%CI[-.375, -.178]), and trial number (mean=-.237, 89%CI[-.331, -.143]). Additionally, EEG decoding performance positively predicted memory (mean=.116, 89%CI[.022, .211]) suggesting that the availability of stimulus information in EEG signals may index an encoding-ready state. Taken together, successful memory ultimately depends on a number of unique factors, including one’s internal state, which is reflected in the reliability of stimulus representations in brain signals.

## Discussion

The current study reveals that the fidelity of neural representations is modulated by both attentional selection (task relevance) and sustained attentional state: visual items are represented more strongly in brain activity when those items are task-relevant and when they are encountered under better, non-lapsing attention. Representational strength of relevant information wanes prior to sustained attentional lapses, supporting the view that poor sustained attention is characterized by decoupling between neural and task processing. Further, irrelevant information processing does not reliably increase prior to lapses, suggesting that poor sustained attentional performance is not simply a failure of selective attention. Finally, we demonstrate a link between attention-related neural processing and memory, with representational fidelity signaling an encoding-ready state. In combination, results provide new insights into the multifaceted nature of attention and its relationship with memory.

Decoding performance of EEG classifiers was reliably modulated by task-relevance. During blocks in which participants made responses based on image category, EEG decoding remained strong for an extended duration relative to blocks in which image category information was task-irrelevant despite similar visual displays in both tasks, suggesting prolonged maintenance of task-relevant image information. This difference emerged following an initial shared peak in decoding performance for both relevant and irrelevant images, indicating that representations of low-level category information may not differ with task-relevance. Additionally, despite low decoding performance overall during Gabor-relevant blocks, orientation decoding was stronger when Gabor patches were task-relevant rather than when they were incidentally-presented. Results align with previous work demonstrating that selective attention may update attentional templates during visual search such that relevant information can be more easily decoded from EEG signals ^3,69^ and extend findings to show that, even when spatial arrangement remains constant, task-relevant information is more-strongly represented in brain signals.

EEG classifiers were trained on data from runs involving passive viewing of stimuli. Despite this, decoding performance when stimuli were task-relevant did not exceed cross-validated performance within localizer (training) runs. From this result, we conclude that selectively attending stimulus information during the CPT did not strengthen category or orientation information in signals, but rather that task-irrelevance led to stronger suppression of category information. However, future work may seek to train a classifier on attended stimuli to test whether selective attention during training impacts generalization across tasks.

On top of task-relevance, we found that dynamic fluctuations in sustained attention influences the time course of decoding. Specifically, we observed worse decoding performance late in the trial for task-relevant images presented prior to sustained attentional lapses. Weak decoding suggests that neural representations of task-relevant images in EEG signals are more degraded under poor sustained attention, in line with hypotheses of decoupling during periods of worse sustained attention ^24,70^. No reliable differences in task-irrelevant decoding suggests that individuals are not simply attending the wrong information prior to lapses. Importantly, we examined decoding accuracy for frequent-category stimuli in the window preceding erroneous responses, such that the weakened decoding indicated a disengaged state that foreshadowed an upcoming sustained attentional lapse. These results suggest that lapses in tasks requiring frequent responses (commission errors) are not purely due to failure of response control or motor inhibition, as has been suggested in previous work ^36^. Rather, while response control likely plays a role in attentional performance, lapses may be jointly shaped by both attention and motor processes ^44^.

Despite our observation that response speed and variance predicted sustained attention performance as measured with behavior (accuracy on infrequent trials), EEG classification results did not support our prediction that decoding performance would differ by response time measures of sustained attention. Response time differences, while predicting lapses behaviorally, were not sufficient to lead to differences in decoding success and did not explain weakened decoding prior to lapses. This finding conceptually replicates recent work suggesting that, despite indexing a more error-prone state, response time measures of attentional state and sustained attentional lapses are characterized by distinct neural mechanisms^16^. Current results suggest that poor decoding performance observed prior to lapses is driven by true differences in brain signals under fluctuating sustained attentional engagement.

Gabor orientation decoding was poor, even when Gabor patches were task-relevant. While expected, as noted in our preregistration, we hypothesized that sustained attention would impact decoding performance. However, we did not observe reliable differences as a function of sustained attention performance for orientation decoding. Poor orientation decoding is likely due to the presentation of Gabor patches in the periphery of the visual field, for which weaker decoding performance has been previously noted ^71^. Future work may seek to test how arrangement of the visual display interacts with selective and sustained attention effects on decoding performance.

Finally, we examined the impact of stimulus-, task-, and attention-related features on memory performance. As expected, previously-reported features like stimulus-specific memorability and lure false-alarmability predicted whether an image would be successfully recognized ^66,67^. Additionally, we found that memory for target images was predicted by classifier performance in the window preceding their presentation, suggesting that EEG classification success, which tracks sustained attention, may reflect a processing state that enables successful mnemonic encoding. Results further support a key link between dynamic attentional and mnemonic states and emphasize the importance of considering attention in models of memory ^46^.

In sum, the current study demonstrates that neural representations are shaped by both attentional prioritization and momentary sustained attention. The fidelity of neural signals was higher for task-relevant items, suggesting that selection strengthens the availability of stimulus information in the brain. Representational strength weakened prior to sustained attentional lapses and also predicted subsequent recognition memory, suggesting that representational strength indexes processing- and encoding-ready states. Results provide novel neural evidence for attentional engagement as well as for a key link between sustained attention and memory.

## Methods

Data was collected as part of a preregistered study (https://osf.io/h9r4e/overview). Deviations from preregistered analyses are noted at the end of the Methods section. Participants were recruited from the University of Chicago and the surrounding community and were compensated for participation with cash, class credit, or a combination of cash and class credit. All aspects of the study were conducted in accordance with protocols approved by the Institutional Review Board at the University of Chicago.

The full experimental procedure included three tasks in total. Task order was always as follows: EEG localizer task, continuous performance task, memory task. Each task is described in detail below.

### EEG localizer task

Participants completed six blocks of the EEG localizer task, named for the fMRI tasks after which it was modeled. Participants were tasked with detecting when the fixation dot presented in the center of the screen changed from red (90% of trials) to green (10% of trials). Additionally, on each trial, a visual stimulus was presented in the background which participants were told was irrelevant for their fixation task. Three blocks, termed “image-category localizer” blocks, included images as the irrelevant stimulus. In the other three blocks, termed “Gabor-orientation localizer” blocks, irrelevant stimuli were Gabor patches with a gray square in their center. Block-specific details are described below. Blocks were presented in an alternating order, with the first block type counterbalanced across participants.

During three image-category localizer blocks, background stimuli were images of food (50%) and vehicles (50%), obtained from the MemCat image dataset ^72^. Images were presented in the center of the screen ∼8.5 cm for 800 ms with a 200 ms inter-trial interval (ITI), 600 trials per block. The fixation dot turned green, requiring participant responses, on 10% of trials, half of which were presented during simultaneous presentation of food images and half during vehicle images. Images were always trial-unique within a block but could be presented up to two times throughout all three image-category localizer blocks. Image presentation order was randomized within blocks.

Gabor-orientation localizer blocks featured background stimuli composed of Gabor patches ∼27.5 cm wide with a ∼8.5 cm gray square in their center. Gabor patches were oriented horizontally on half of trials and vertically on the other half of trials, presented in a random order. Patches were presented for 800 ms with a 200 ms ITI, 600 trials per block. The fixation dot changed from red to green on 10% of trials, half of which included horizontally-oriented Gabor patches and half of which included vertically-oriented Gabor patches.

Participants were able to rest between blocks. For both image-category and Gabor-orientation localizer tasks, fixation color change detection was performed as an attention check task. Trials on which the dot turned green were excluded from EEG analyses.

### Continuous performance task

Following the EEG localizer task, participants performed four blocks of a continuous performance task (CPT) designed to test fluctuations in sustained attention. This task was a modified version of a not-X CPT during which participants were instructed to respond with a button press to frequent-category stimuli and withhold their button to rare targets ^21^. Stimuli in the current study always comprised an image (∼8.5 cm) superimposed in the center of a Gabor patch (∼27.5 cm). Stimuli were presented for 800 ms, followed by a 500 ms blink period during which participants were instructed to blink—to reduce the number of blink artifacts during trials—and lastly a 200 ms ITI. Each block included 600 trials total. During two blocks, termed “Image-relevant CPT” blocks, participants were instructed to make a judgement about the category of the image and Gabor patches were task-irrelevant. During the other two blocks, “Gabor-relevant CPT” blocks, participants made responses based on the orientation of the Gabor patches and images were task-irrelevant. Task details for each block are described below. All images presented during the CPT were trial-unique and were not presented in any other localizer or CPT block. Image- and Gabor-relevant blocks were performed in alternation, with the first block counterbalanced across participants and unique from the order of the localizer task.

During image-relevant CPT blocks, images were drawn from one of three image categories. Two were frequent categories—food images (45% of trials) and vehicle images (45%). The third category—animal images (10% of trials)---served as rare targets. Participants were instructed at the beginning of the block to respond by pressing the space bar when they saw images of food or vehicles, and to withhold their response when the image was an animal image. Responses to frequent-category images occurring before the onset of the subsequent trial were considered correct. Task-irrelevant Gabor patches were oriented either horizontally (50% of trials) or vertically (50% of trials).

Gabor patches during Gabor-relevant CPT blocks were oriented in one of three directions. Frequently, patches were oriented vertically (45% of trials) or horizontally (45% of trials). On rare target trials (10%), patches were oriented diagonally with a 45 degree left tilt. Participants were instructed to respond by pressing the space bar when patches were oriented horizontally and vertically but withhold their response when the patch was oriented diagonally. Responses to horizontal and vertical Gabor patches occurring before subsequent stimulus onset were considered correct. Task-irrelevant images were either food images (50% of trials) or vehicle images (50% of trials).

### Two-alternative forced choice memory task

Finally, participants completed a surprise two-alternative forced choice (2AFC) memory task for image stimuli presented during CPT blocks. On each trial, one old and one new image were presented to the left and right of fixation, presentation order counterbalanced across trials. Participants were instructed to report the identity of the old image and provided a confidence rating—definitely the image on the left, maybe the image on the left, maybe the image on the right, definitely the image on the right—using the z, x, n, and m keys, respectively. The new image was always a lure from the same image category. Task instructions remained on the screen for all trials and trials were self-paced such that participants could take as long to respond as needed. Memory was tested for 60 relevant food images, 60 relevant vehicle images, 60 relevant animal images, 60 irrelevant food images, and 60 irrelevant vehicle images, resulting in a total of 300 memory trials. Relevant image memory was always tested first, followed by irrelevant image memory. EEG data from the memory task is not analyzed as participants were not required to use the chin rest during this task. One participant chose to leave prior to the memory task. Therefore, memory analyses include data from 24 participants.

### EEG data collection

EEG data was collected using 30 active electrodes (actiCHamp Plus, Brain Products, Munich, Germany) within an electrically shielded Faraday cage. Electrodes were placed at the following sites: Fp1, Fp2, F7, F3, Fz, F4, F8, FT9, FC5, FC1, FC2, FC6, FT10, T7, C3, Cz, C4, T8, CP5, CP1, CP2, CP6, P7, P3, Pz, P4, P8, O1, Oz, O2. The ground electrode was placed at Fpz. Two electrodes were placed on the right and left mastoids. During data collection, electrodes were referenced to the right mastoid. Offline, data were re-referenced to the algebraic average of the left and right mastoids. Data were filtered online with a high cutoff of 140 Hz and digitized at a sampling rate of 500 Hz using BrainVision Recorder (Brain Products, Munich, Germany) running on a PC.

Electrooculogram (EOG) activity was recorded using passive electrodes. Horizontal EOG activity was recorded using electrodes placed approximately 1cm from the outer canthus of each eye. Vertical EOG activity was recorded using a pair of electrodes placed above and below the right eye. A ground electrode was placed on the left cheek.

### Eye tracking

Gaze position and pupil size were collected using a desk-mounted infrared eye-tracking system (EyeLink 1000 Plus, SR Research, Ottawa, Ontario, Canada) sampled at 1000 Hz. A chin rest helped participants maintain a stable head position. The eye tracker was calibrated at the start of each EEG localizer and CPT block.

### Preprocessing and artifact rejection

EEG data were bandpass filtered between 0 and 80 Hz and epoched from 150 ms prior to trial onset to 800 ms after trial onset. Baseline correction was applied using the time period -150 to 0 ms prior to stimulus onset.

Artifact rejection was performed using an automated detection procedure on EEG and EOG data. Trials containing artifacts were excluded from EEG analyses but retained for behavioral analyses. Due to participant motion over long task blocks, gaze position and pupil data was lost for large portions of task blocks making it unsuitable for artifact rejection. However, gaze position and pupil data were retained for control analyses.

Trials containing saccades or blinks were rejected if any of the following criteria were met: 1) if the difference in the mean activity in the first and second halves of an 80 ms sliding window (advanced in 10 ms increments) exceeded 60 microvolts in EEG channels or 50 microvolts in EOG channels; 2) if the range of activity within a 200 ms window exceeded 100 microvolts in EEG channels or 300 microvolts in EOG channels, or 3) if the absolute value of channel activity exceeded a difference from baseline of 100 microvolts in EEG channels or 300 microvolts in EOG channels. Trials containing linear drifts were excluded if the slope of a linear fit across the epoch exceeded 75 microvolts and the coefficient of determination (R^2^) exceeded 0.3. Finally, trials with dropout or in which any channel flatlined for at least 200 ms were excluded.

Known noisy electrodes, frontal EEG channels Fp1 and Fp2, were excluded from preprocessing and subsequent analyses for all participants. In the case that noise in a single additional EEG channel resulted in excessive artifacts, that channel was excluded from preprocessing and all subsequent analyses. On average, 26.9% of trials were excluded after preprocessing (SD=15.0%).

### Participant exclusion

Participants were excluded from analyses if they met any of the following criteria, established in the preregistration: 1) Average behavioral performance (sensitivity, *A’*) during CPT blocks fell below 2.5 SD of the group average, or 2) More than half of CPT trials were excluded due to artifact rejection. No participants were excluded for low CPT performance. Four participants were excluded for excessive artifact rejection. The final sample size included 25 participants (mean age=22.24 years, SD=2.93 years). The gender breakdown was as follows: 15 men, 9 women, 1 non-binary person.

### Behavioral methods

Individual-level behavioral performance on the CPT and during the localizer was quantified using *A’*, a non-parametric measure of sensitivity. The formula for calculating *A’* is as follows, where *hits* represents the hit rate and *FAs* represents false alarm rate:

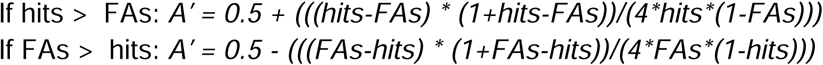

Sensitivity *(A’)* was calculated for each block separately and averaged within participants across image-relevant and Gabor-relevant blocks to obtain a measure of performance per block type.

Frequent responding during the CPT provides a behavioral index of sustained attentional state throughout the duration of the task. Extensive previous work has linked momentary measures of response time speed and variance to sustained attention performance ^21,22,57^. Specifically, periods of slow ^53,55^ and consistent ^54^ responding are more likely to precede successful withholding of presses to rare targets. Further, these measures uniquely predict upcoming lapses ^48,49^ suggesting that they index separate components of attention-related behavior.

To obtain a continuous measure of response speed, time courses of responses to frequent, correct responses were linearly detrended within block to account for speeding over time and z-scored within block. To calculate a measure of response time variance, we computed the absolute value of detrended and z-scored response time courses to obtain a measure of deviation from the average response time. Finally, missing values were linearly interpolated from surrounding trials.

We aimed to test whether response speed and variance uniquely predicted upcoming sustained attentional lapses, i.e., presses to infrequent target stimuli, replicating previous work ^48,49^. We calculated the average response speed and variance in the three-trial window preceding an infrequent target. Using a Bayesian logistic mixed effects modeling approach, we fit models predicting accuracy on infrequent trials from pre-trial response speed, variance, their interaction, and a categorical predictor of CPT block type. Finally, a subject-level random intercept was included. Models were fit using the package *brms* in R using default priors (4 chains, 10,000 iterations each).

Individual-level memory performance was calculated as sensitivity (*A’*) across memory trials for relevant and irrelevant images separately. Images were considered correctly-remembered if participants reported a confident, accurate judgement. Following our preregistration, we also report memory accuracy for both confident and less confident memory judgements. When calculating memory performance for confident judgements, hit rates were calculated as the number of old images confidently remembered divided by the total number of confident reports (hits and false alarms). False alarm rate was calculated as the number of confident false alarms divided by the total number of confident reports. Due to the two-alternative forced choice nature of the task, hit rates and false alarm rates always summed to 1.

We sought to examine what features influenced successful memory judgements in the current study. Primarily, we were interested in whether measures related to selective attention (task-relevance), salience (stimulus frequency), and sustained attention (pre-trial RT speed and variance) predicted memory when controlling for known predictors of memory such as stimulus memorability ^66^, the likelihood of the paired memory lure to be incorrectly reported as remembered (false-alarmability ^67^), and trial number. Memorability scores for all images in the MemCat dataset were provided by the authors ^72^. For the current analyses we used memorability without correction for false alarms. False-alarmability was calculated for memory lure images as the number of false alarms divided by the total number of presentations in the original dataset.

To test the influence of these features on memory we fit Bayesian logistic mixed effects models predicting memory accuracy from effects of task-relevance (i.e., whether an image was presented in an image-relevant or a Gabor-relevant CPT block), category frequency (i.e., whether an image was a rare animal image), memory trial number, stimulus memorability, memory lure false-alarmability, and pre-trial speed, variance, and their interaction. We additionally included a subject-level random intercept term.

### EEG Classifiers

The primary aim of this study was to test the impacts of attention on the representational strength of stimulus information as measured by EEG decoding performance. We hypothesized that both selective attention—i.e., whether a stimulus was task-relevant—and sustained attentional state—i.e., whether a stimulus was presented during a period of better sustained attention—would result in stronger EEG decoding performance as the result of more actively-maintained stimulus information in mind.

We trained within-subject EEG linear discriminant analysis (LDA) classifiers to decode image category (food vs. vehicles) or Gabor orientation (horizontal vs. vertical) on all EEG localizer block trials. Classifiers were trained on the raw EEG activity across channels at every time point for all trials in localizer blocks. Training on these blocks ensured that data were completely independent from testing CPT blocks and removed any confound of responding on classifier training. Trained models were tested on CPT block trials.

We calculated the average performance of classifiers as a function of selective attention by splitting trials into whether the classifier was predicting the category of task-relevant or task-irrelevant stimuli. To test whether pre-trial stimulus representations predicted upcoming attentional performance, we examined classifier performance for the three trials immediately preceding an attentional lapse—i.e., an incorrect press to an infrequent target—versus those preceding a correctly-withheld response. Lastly, to test the impacts of sustained attentional state, trials were split by whether they appeared in the highest quartile (in-the-zone state) or lowest quartile (out-of-the-zone state) of sustained attention using a quartile split on the response speed and variance time courses.

Primary analyses tested whether the time course of EEG decoding performance meaningfully differed as a function of attention when time-locking to the onset of the visual stimulus. To do so, we tested classifier performance at the same time point in the trial at which the classifier was trained. Performance for all EEG decoders was quantified using the area under the curve. This analysis allows us to test whether the time course of stimulus decoding across the trial differs as a function of attention. Differences between time courses across subjects were quantified with Bayes Factors at every time point ^73^. Bayes Factors were calculated using a paired test with a default Cauchy prior with a medium scale parameter (r∼.707).

For comparison, we also performed k-fold cross-validated classification for both images and Gabor patches during the localizer task blocks. The EEG decoder was trained on two-thirds of localizer trials and performance was tested on the left-out third of trials. This process was repeated until each third of trials served as the held-out set. Performance was averaged across the three folds to return the average decoding performance for each subject.

### Event-related potentials

To examine whether activity in response to the presentation of frequent stimuli differed under different sustained attentional states, we calculated the average activity in trials presented under good sustained attention (the three-trial window preceding a correctly-withheld response to an infrequent target) and poor sustained attention (the window preceding a commission error or lapse in response to an infrequent target) in each channel across the scalp. We then calculated the difference between these average activity time courses for each subject.

### Time-frequency analysis

We performed a time-frequency decomposition analysis to examine differences in oscillatory power under different sustained attentional states. Specifically, we examined trials in the window preceding an infrequent target as a function of performance on that target trial. Using a set of posterior electrodes, P3/P4/Pz/O1/O2/Oz, we convolved EEG time series with a Morlet wave for frequencies between 2 and 30 Hz in 1 Hz steps. The number of cycles increased based on frequency using the calculation n_cycles_=frequency/3, resulting in wavelets ranging from approximately 1 cycle at 3 Hz to 10 cycles at 30 Hz. This analysis was implemented using the *tfr_array_morlet* function for MNE in python. For each subject, we averaged the resulting power matrix across trials as a function of sustained attention and calculated the difference in the time-frequency matrices. The resulting matrix demonstrates power differences in frequency bands between good and poor sustained attentional states.

### Decoding from eye data

To examine whether any differences in EEG decoder performance may be driven by artifacts due to eye gaze or pupil size, we also trained LDA classifiers using the data collected by the infrared eye tracker (gaze position, pupil size) and raw activity from vertical and horizontal EOGs. Before training, missing values in eye data were replaced with the average feature value across all available EEG localizer trials (training dataset) or CPT trials (testing dataset). Eye classifiers were trained and tested identically to EEG classifiers to determine whether decoding time courses show similar modulation by attention. We compared predictions from classifiers trained on EEG and eye data by correlating trial-wise predictions at each time point.

### Predicting memory from EEG decoding

Previous work suggests that memory for infrequent targets is predicted by the attentional state in which they are encountered, as measured behaviorally ^48,49,53–55^ and neurally ^14,74^. Here, we examined whether EEG decoding performance, a hypothesized index of attentional state, predicts subsequent memory for infrequent target images. We tested this possibility by fitting Bayesian logistic mixed effects models predicting infrequent trial memory success from EEG classification confidence—that is, the probability that a food image was correctly classified as food, or a vehicle image as a vehicle—in the three trials preceding its presentation during the CPT. Decoder confidence was averaged across the entire trial duration for each image. We additionally included variables expected to be related to image memory performance as controls: image memorability, paired lure false-alarmability, and memory trial number, as well as a random intercept of subject. Image memorability values were obtained from the shared MemCat image database (memorability without false alarm correction). We calculated paired lure false-alarmability using data shared in the MemCat image database. Specifically, false-alarmability was calculated as the number of false alarms made to a given image divided by the total number of times that image was tested.

### Differences from preregistration

For transparency, we note changes from preregistered analyses here. First, Bayesian statistics are used in lieu of parametric significance testing when evaluating differences in decoding time courses throughout the manuscript.

We initially preregistered indexing sustained attentional state using a median split on both response speed and variance time courses. The decision to instead use upper and lower quartiles of both response speed and variance was made to improve power by isolating the most extreme sustained attentional states. However, results and conclusions do not differ between these analysis choices.

To better explore differences in neural representational strength under different sustained attentional states, we analysed decoding performance in the three-trial window prior to infrequent trials as an exploratory analysis. While rare, infrequent trials provide a more objective measure of sustained attentional state than response time indices. Capturing objective lapses in sustained attention enabled the differences observed in decoding accuracy in the preceding three-trial window.

Finally, one preregistered exploratory analysis proposed testing whether oscillatory power predicted successful memory for stimuli. However, we decided that a more relevant analysis would test differences in oscillatory power under better and worse sustained attention conditions, as the primary aim of this study was to examine neural signatures of attention. This analysis is reported in the main text.

## Acknowledgements

The authors would like to thank Dr. John Veillette for his help initializing the EEG and eye tracking systems for data collection and Dr. Henry Jones for thoughtful conversations on methods and analyses.

