## Supplementary materials for "Degraded neural representations foreshadow failures of sustained attention"

**Supporting Information**

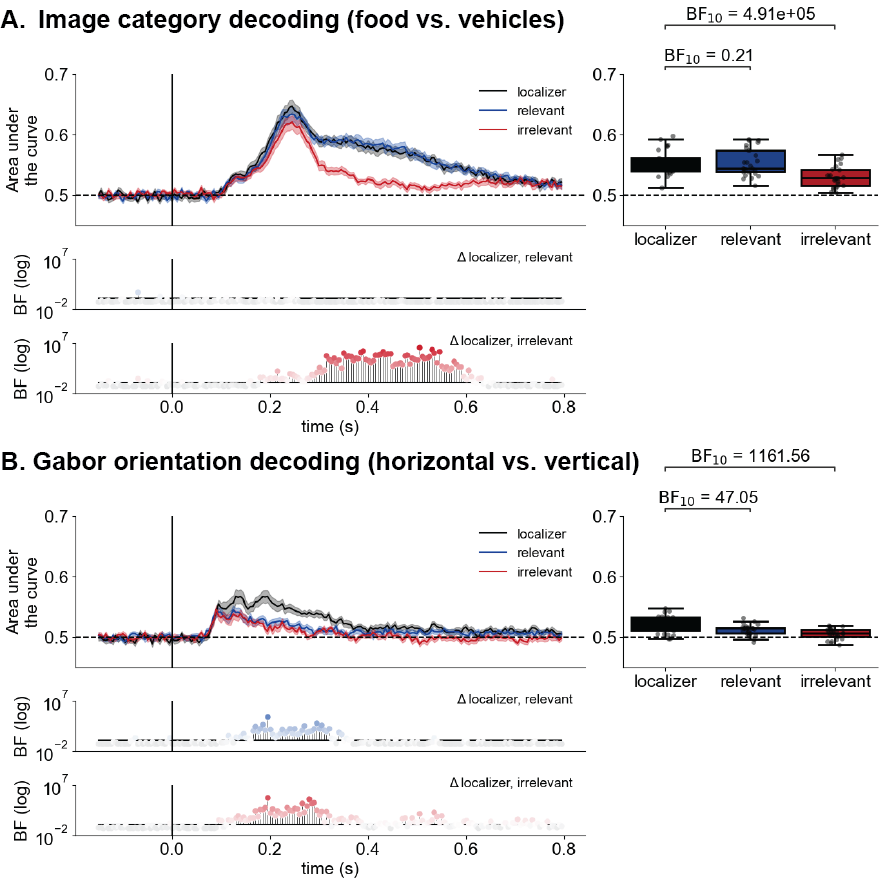

*Supplementary Figure 1. Relevant decoding does not exceed cross-validated decoding time courses.* We compared the time courses trained and tested during the localizer task, during which participants passively viewed stimuli, to those observed during the CPT. (A) When decoding image categories, relevant image decoding performance was very similar to the cross-validated localizer time course, whereas irrelevant image decoding was decreased. (B) When decoding Gabor orientations, the cross-validated localizer time course showed higher and more extended performance than during either Gabor-relevant or Gabor-irrelevant CPT blocks.

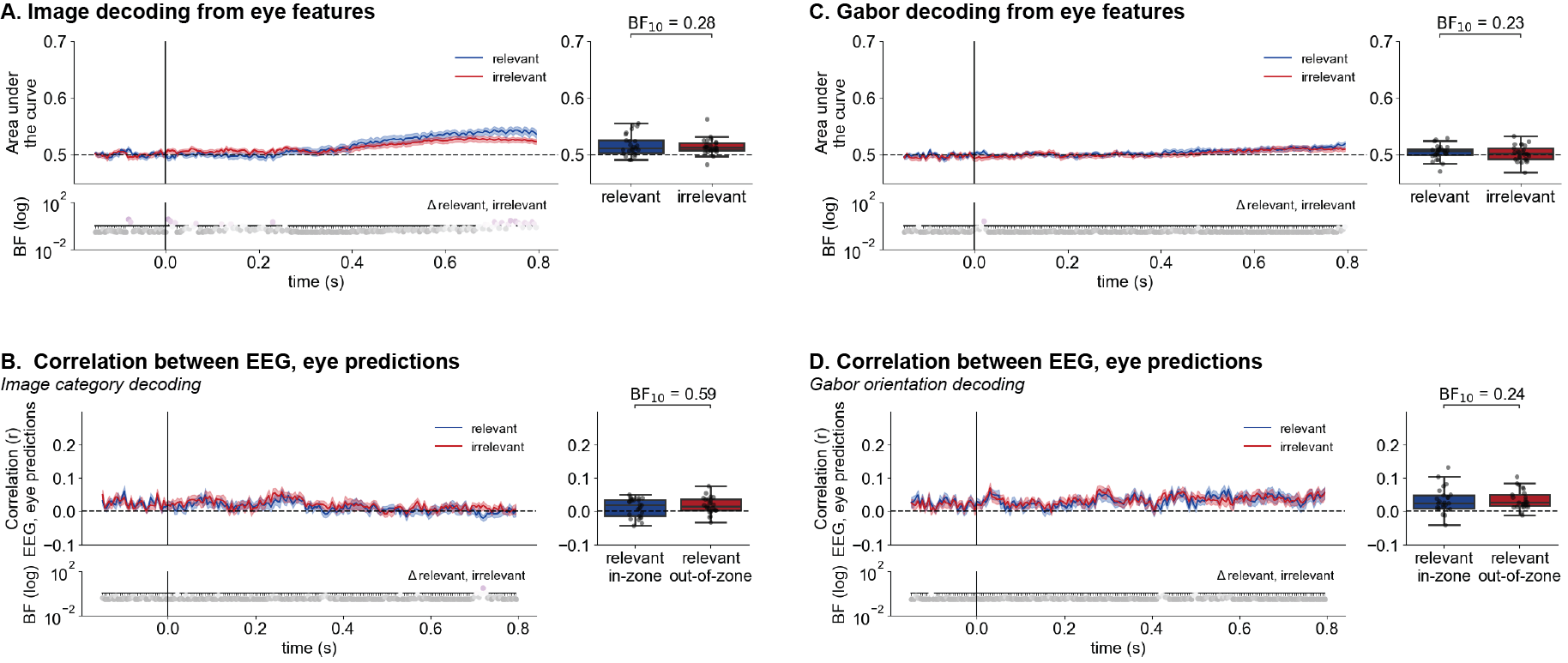

*Supplementary Figure 2. Decoding from eye features is distinct from EEG.* Classifiers using eye features resulted in distinct decoding time courses for (A) image category and (C) Gabor orientation relative to those trained on EEG signals. Further, predictions from EEG-based and eye-based classifiers were minimally correlated across trials for (B) images and (D) Gabors, and this correlation did not differ between task-relevant and task-irrelevant predictions. This suggests that differences in decoding accuracy observed as a function of attentional selection (task relevance) are not driven by differential dependence on eye-related features.

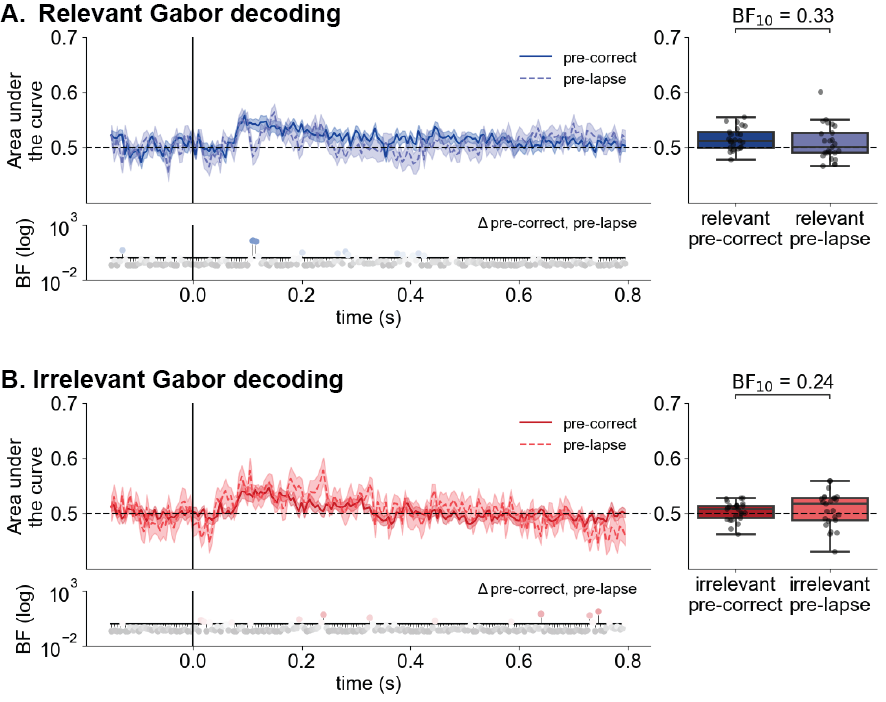

*Supplementary Figure 3.* *Gabor decoding does not differ preceding sustained attentional lapses.* Decoding performance for (A) relevant and (B) irrelevant Gabor orientations showed only small differences as a function of sustained attentional state. Trials under good sustained attention states were those preceding correctly withheld responses to infrequent target stimuli (pre-correct), while trials preceding lapses (pre-lapse) were considered to be in a poor sustained attentional state. We did not observe differences when averaging performance across the trial (0-800ms), as shown in the box-and-whisker plots.

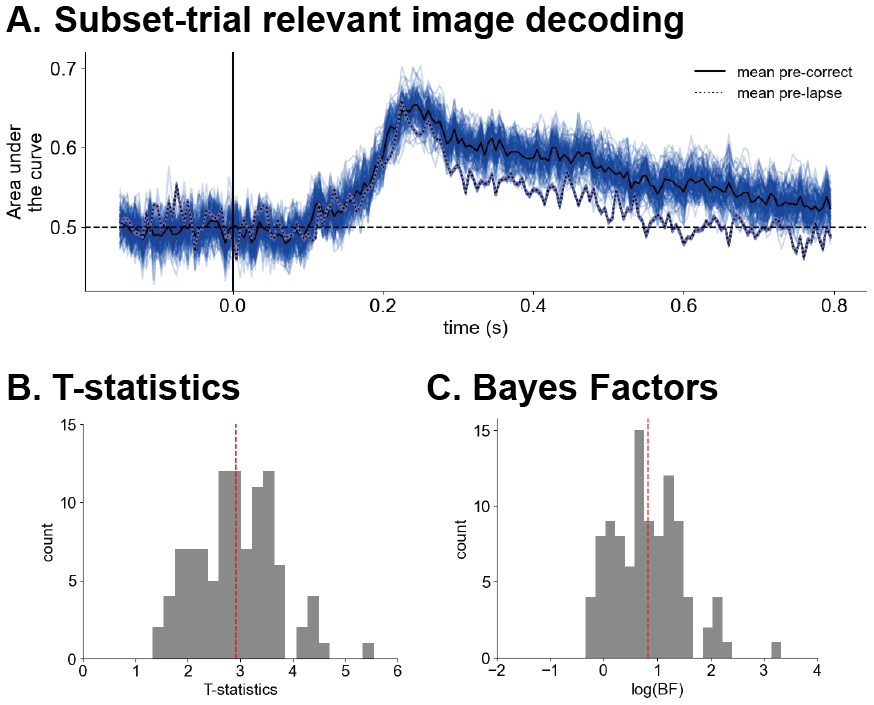

*Supplementary Figure 4. Weakened pre-lapse decoding is not explained by smaller trial counts.* To ensure that differences in trial count between pre-correct and pre-lapse trials did not explain differences in decoding performance between conditions, we performed a subsetting analysis. For each participant, the number of trials used for testing both the pre-correct and pre-lapse decoding was subset to the smaller of the two conditions. A random set of trials from the larger condition was used for classifier testing. This was repeated 100 times with different subsets of the larger condition. (A) Pre-correct decoding remained more accurate than pre-lapse decoding when trial counts were equated. The distribution of (B) positive T-statistics and (C) Bayes Factors from paired t-tests comparing the mean pre-correct and pre-lapse decoding performance over the trial reveal that pre-correct decoding was higher than pre-lapse decoding across iterations. Red dashed lines reflect the mean.

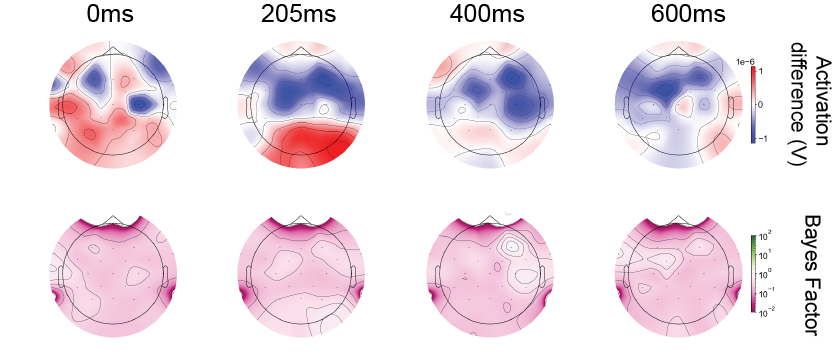

Supplementary Figure 5. *Minimal differences in event related potentials as a function of sustained attention.* Topographic maps reflect the difference in average activation across the scalp as a function of sustained attention (trials preceding correctly-withheld responses minus trials preceding lapses) in frequent images presented in the three trials preceding an infrequent target. Bayes factors reflect evidence of a reliable sustained attention-related activation difference at every channel. Bayes factors greater than 1 are plotted in green and reflect evidence of an effect.

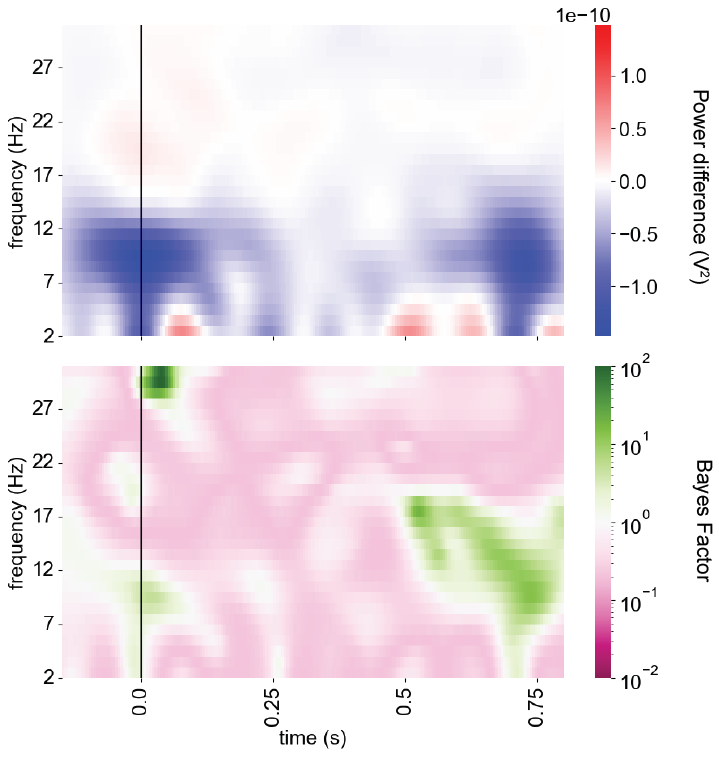

*Supplementary Figure 6. Time-frequency decomposition for good vs. poor sustained attentional states.* Differences in time-frequency activity in posterior electrodes (pre-correct minus pre-lapse trials) reveal greater alpha (8-12 Hz) power at stimulus onset (time=0 s) and prior to stimulus offset (time=0.8 s) prior to sustained attentional lapses. Bayes factors reflect evidence of an effect (distribution reliably different from 0) at every frequency and time point. Bayes factors greater than 1 are plotted in green and reflect evidence of an effect.

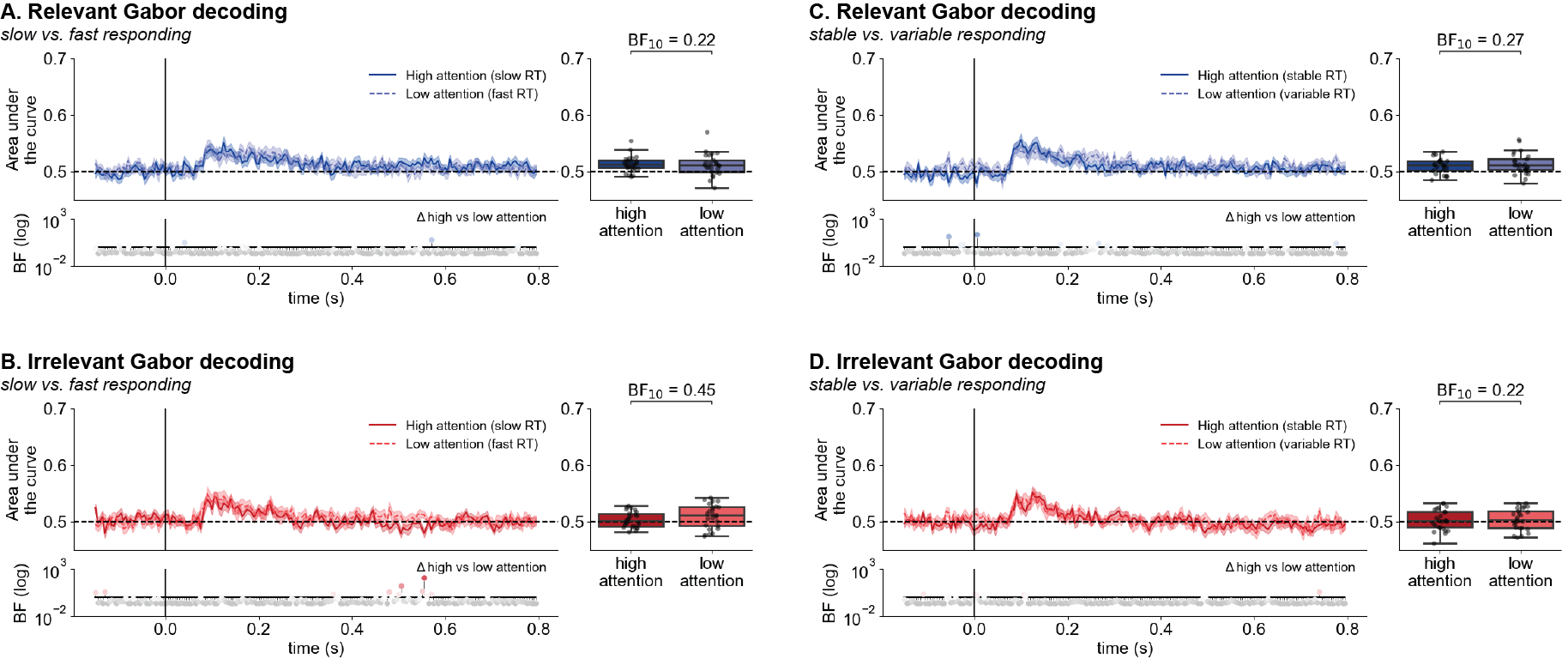

*Supplementary Figure 7. Minimal effect of response time-indexed sustained attentional state on Gabor orientation decoding.* We observed only small, transient differences in decoding performance for Gabor orientation as a function of response-indexed sustained attentional state using response time (RT) speed (A-B) and variance (C-D). When averaged across the trial, no differences were observed as a function of sustained attention. Box-and-whisker plots reflect subject-wise mean area under the curve across the trial (0-800 ms after onset).

| **Predictor** | **Mean estimated effect** | **89% Credible Interval** |
| --- | --- | --- |
| Memorability | .131 | [.093, .170] |
| Paired lure false-alarm-ability | -.283 | [-.322, -.243] |
| Memory trial number | -.027 | [-.102, .047] |
| Relevance (relevant vs. irrelevant) | .231 | [.075, .386] |
| Frequency (infrequent vs. frequent) | .663 | [.550, .776] |
| Preceding RT speed | .091 | [.039, .143] |
| Preceding RT variance | .025 | [-.020, .073] |
| RT speed:RT variance | -.020 | [-.040, 4.38*10^-4^] |

*Supplementary Table 1.* *Effects predicting both confident and non-confident memory judgements.* As a preregistered analysis, we tested what features predicted successful selection of the old stimulus during a two-alternative forced-choice memory task regardless of confidence rating. We fit a Bayesian mixed effects logistic model and report the mean estimated effect and 89% credible interval here. Credible intervals that do not include 0 are considered evidence of an effect. We observe that image memorability, the likelihood of the paired lure stimulus to elicit false alarms (paired lure false-alarmability), and stimulus frequency remain strong predictors of memory. Additionally, task-relevance and response speed emerge as predictors of memory that did not previously predict confident memory judgments alone.

| **Predictor** | **Mean estimated effect** | **89% Credible Interval** |
| --- | --- | --- |
| Memorability | .085 | [-.012, .185] |
| Paired lure false-alarm-ability | -.426 | [-.525, -.327] |
| Memory trial number | -1.11*10^-3^ | [-.100, .100] |
| Pre-trial EEG decoding performance | -.038 | [-.137, .061] |

*Supplementary Table 2.* *Effects predicting both confident and non-confident memory for infrequent target images.* We fit a Bayesian mixed effects logistic model predicting memory for rare target images where successful identification of the old stimulus is considered correct regardless of confidence. We report the mean estimated effect and 89% credible interval here. Credible intervals that do not include 0 are considered evidence of an effect. Only the likelihood of the paired lure image to elicit false alarms (paired lure false-alarm-ability) predicted image memory in this model.
